# Kernel-based genetic association analysis for microbiome phenotypes identifies host genetic drivers of beta-diversity

**DOI:** 10.1101/2021.10.15.464608

**Authors:** Hongjiao Liu, Wodan Ling, Xing Hua, Jee-Young Moon, Jessica S. Williams-Nguyen, Xiang Zhan, Anna M. Plantinga, Ni Zhao, Angela Zhang, Rob Knight, Qibin Qi, Robert D. Burk, Robert C. Kaplan, Michael C. Wu

**Affiliations:** Department of Biostatistics, University of Washington, Seattle, WA; Public Heath Sciences Division, Fred Hutchinson Cancer Research Center, Seattle, WA; Department of Epidemiology and Population Health, Albert Einstein College of Medicine, New York, NY; Department of Biostatistics and Beijing International Center for Mathematical Research, Peking University, Beijing, China; Department of Mathematics and Statistics, Williams College, Williamstown, MA; Department of Biostatistics, Johns Hopkins University, Baltimore, MD; Department of Pediatrics, University of California, San Diego, La Jolla, CA

## Abstract

Understanding human genetic influences on the gut microbiota helps elucidate the mechanisms by which genetics affects health outcomes. We propose a novel approach, the covariate-adjusted kernel RV (KRV) framework, to map genetic variants associated with microbiome beta-diversity, which focuses on overall shifts in the microbiota. The proposed KRV framework improves statistical power by capturing intrinsic structure within the genetic and microbiome data while reducing the multiple-testing burden. We apply the covariate-adjusted KRV test to the Hispanic Community Health Study/Study of Latinos in a genome-wide association analysis (first gene-level, then variant-level) for microbiome beta-diversity. We have identified an immunity-related gene, *IL23R*, reported in previous association studies and discovered 3 other novel genes, 2 of which are involved in immune functions or autoimmune disorders. Our findings highlight the value of the KRV as a powerful microbiome GWAS approach and support an important role of immunity-related genes in shaping the gut microbiome composition.

## Introduction

The human microbiome plays an important role in host health and is involved in fundamental body functions such as metabolism and immune response [1, 2]. While environmental factors have a large influence on microbiome composition [3], it is still of interest to study the effect of human genetic variation on the microbiome: such studies provide clues as to the biological mechanisms by which genetics may influence health outcomes. As a notable example, elevated abundance of *Bifidobacterium*, a genus of beneficial gut bacteria that utilizes lactose as an energy source, has been associated with a non-persistence genotype of the human lactase gene (*LCT*), which typically results in lactose intolerance [4, 5, 6]. Such an association implies a potential mediating role of the microbiome in the relationship between host genetics and metabolic outcomes, where the presence of Bifidobacteria may provide some level of lactose tolerance to lactase non-persistent individuals [4].

Many studies have sought to identify genetic variants that influence microbial composition, and most of them incorporate microbiome characteristics as phenotypes in genome-wide association studies (GWAS). Typical analyses marginally test the association between abundances of individual taxa and genotypes of individual genetic variants [5, 7, 8, 9]. Such analyses often suffer from a low statistical power, due to a large multiple-testing burden and failure to accommodate inherent structure in microbiome and genetic data, e.g., phylogenetic relationships among taxa and epistasis among genetic variants.

As the microbiome functions as a community, an alternative microbiome phenotype is beta-diversity, the dissimilarity in overall microbiome profiles between individuals. Beta-diversity analysis represents a standard mode of analysis in microbiome profiling studies as it focuses on discovery of concerted shifts in the community rather than individual taxa. However, few studies have considered beta-diversity as a trait of interest in microbiome GWAS and there is no standard strategy. Some studies [6, 10] have performed principal coordinates analysis (PCoA) on the pairwise beta-diversity matrix and evaluated the association between the top principal coordinates (PCo’s) and the genotype of each genetic variant. Such a strategy could suffer from power loss, as the top PCo’s may not fully capture the variation within the microbiome data. Hua et al. [11] assumed a linear model between the pairwise beta-diversity and the pairwise genetic distance at each genetic variant and developed a score test in a tool called microbiomeGWAS. Rühlemann et al. [12] adopted a distance-based multivariate analysis of variance (MANOVA) approach called distance-based F test [13] and evaluated the difference in beta-diversity among the different genotype groups for each genetic variant. These approaches still test one variant at a time and are subject to a stringent genome-wide significance threshold. Studies using the above approaches have identified loci within genes involved in immunity [6, 12], vitamin metabolism [10] and complex diseases such as type 2 diabetes [14]. In our study, we aim to further improve statistical power with a novel approach and bring more discoveries from microbiome GWAS.

Here we propose to assess the association between groups of variants at the gene level and the overall microbiome composition, characterized by beta-diversity, at the community level. Community-level analyses and multi-variant testing have been shown to be powerful in microbiome [15, 16] and genetic studies [17], respectively, due to their ability to capture innate structure and correlation within the data, while reducing the multiple-testing burden. Using the recently developed kernel RV (KRV) framework [18, 19], we summarize the individuals’ microbiome (or genetic) characteristics by a pairwise similarity matrix called “kernel” matrix, where each entry in the matrix represents similarity in microbiome (or genetic) profiles between a pair of individuals. Microbiome similarity can be obtained by transforming known beta-diversity measures, while genetic similarity can also be characterized in various ways, such as the average genotype matching over all genetic variants. The association between microbes and genetics is then assessed via comparing similarity in microbiome profiles to similarity in genetic profiles across all pairs of individuals. Intuitively, if the genetics is associated with the microbiome, we would expect the pairwise microbiome profiles to be similar whenever the pairwise genetic profiles are similar. In particular, the test statistic is the normalized Frobenius inner product, a measure of correlation, between the two kernel matrices.

Although the KRV is a potentially powerful approach for microbiome GWAS, due to the nature of microbiome kernels, the KRV framework has difficulties in controlling for covariates such as population structure, which is imperative for any genetic association analysis. Here we extend the original KRV framework to allow for flexible covariate adjustment.

We apply the covariate-adjusted KRV to the Hispanic Community Health Study/Study of Latinos (HCHS/SOL) [20, 21] via a two-stage (first gene-level, then variant-level) genome-wide association analysis. This is the first study to investigate the genetic effect on the overall gut microbiome composition, characterized by beta-diversity, in Hispanic/Latino populations. We have identified a gene (*IL23R*) reported in a previous microbiome genetic association study and discovered other novel genes related to immune functions. Furthermore, we have identified 311 significant variants within these genes. In addition, our simulation results show that the covariate-adjusted KRV maintains valid type I error rates in the presence of confounding and has a much greater power than other single-trait-based competing methods across a range of scenarios. Together, our proposed approach demonstrates good statistical properties and provides a powerful way to study the effect of human genetic variation on microbiome composition.

## Results

### Overview of covariate-adjusted KRV

We aim to assess the covariate-adjusted association between genotypes of multiple genetic variants within a gene and abundances of multiple microbiome taxa, using the previously developed KRV framework. We now give an overview of the original KRV framework and extend it to allow for covariate adjustment.

The KRV framework has been proposed by Zhan et al. [18, 19] to evaluate the general association between a group of genetic variants, *G*, and a group of traits, *Y*. Suppose we have genotype data of *m* genetic variants and phenotype data of *q* traits available for *n* unrelated individuals. For the *i*th subject, let ***g***_*i*_ = (*g*_*i*1_, *· · ·, g*_*im*_)^*T*^ be the set of genotypes, where *g*_*il*_ ∈ {0, 1, 2} represents the number of minor alleles for the *l*th variant; let ***y***_*i*_ = (*y*_*i*1_, *· · ·, y*_*iq*_)^*T*^ be the set of traits. Example phenotypes in previous studies include expression values of multiple genes from a particular pathway [18] and levels of multiple amino acids [22]. In the context of microbiome GWAS, we treat the microbiome as the phenotype. Specifically, ***g***_*i*_ represents the genotypes of *m* genetic variants within a particular gene, and ***y***_*i*_ represents the abundances of *q* microbiome taxa that form the microbiota.

Let *k*(***g***_*i*_, ***g***_*j*_) be a kernel function that measures the similarity in genetic profiles between individual *i* and *j*. Let *𝓁* (***y***_*i*_, ***y***_*j*_) be another kernel function that measures the similarity in phenotypic profiles between *i* and *j*. Specific choices of kernel functions in the context of microbiome GWAS are discussed in Choice of kernels. We can then define a kernel matrix ***K*** *∈* ℝ^*n×n*^, where the (*i, j*)-th entry of ***K*** is *k*(***g***_*i*_, ***g***_*j*_). Similarly, we define another kernel matrix ***L*** *∈* ℝ^*n×n*^ such that ***L***_*ij*_ := *𝓁* (***y***_*i*_, ***y***_*j*_). The matrices ***K*** and ***L*** can be viewed as pairwise similarity matrices for genotypes and phenotypes, respectively. We further center the two kernel matrices: let 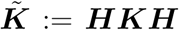 and 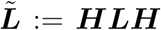, where ***H*** = ***I*** *−* **11**^*T*^ */n* is a column-centering matrix. Then the KRV coefficient that evaluates the relationship between the genetic variants and the traits is defined as

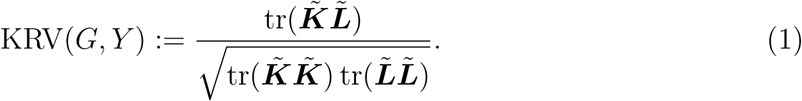

Intuitively, the KRV coefficient compares genotypic similarity to phenotypic similarity across all pairs of individuals. A large KRV coefficient indicates that the pairwise similarity pattern in genetic profiles well resembles the pairwise similarity pattern in phenotypic profiles, which implies that the genetic variants are associated with the traits in a certain way. On the other hand, the KRV coefficient can also be viewed as a multivariate and non-linear extension of the Pearson’s sample correlation coefficient *r*: when both ***g***_*i*_ and ***y***_*i*_ are one-dimensional and we use the linear kernel functions 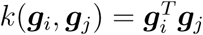 and 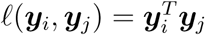, the KRV coefficient is exactly equivalent to *r*^2^. To perform hypothesis testing, the permutation distribution of the KRV statistic under the null hypothesis of no association between genetics and phenotypes can be approximated by a Pearson Type III distribution [18], allowing us to obtain a p-value and assess the significance of the association at a given significance level.

The above framework does not take into account any covariates that might be involved in a typical genetic association study. Unaccounted confounders, such as population structure, can lead to spurious associations in GWAS studies [23]. Now suppose that, for each individual *i*, we have a set of covariates ***x***_*i*_ = (1, *x*_*i*1_, *· · ·, x*_*ip*_)^*T*^ *∈* ℝ^*p*+1^; let ***X*** *∈* ℝ^*n×*(*p*+1)^ be the sample covariates matrix such that the *i*-th row of ***X*** is 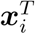. Assume that ***X*** has full rank. We intend to assess the association between the genetic variants and the phentoypes, after adjusting for the effects of the covariates ***X***. Previous studies, including the original KRV framework, have suggested using a residual-based approach [17, 24, 18], where we first regress out the covariates from each raw phenotype and then construct the phenotype kernel matrix using the resulting residuals. Such an approach is not feasible for microbiome data, as popular microbiome kernels (e.g., the Bray-Curtis kernel and the weighted UniFrac kernel) require the input to be discrete taxa count data, which is not satisfied by the covariate-adjusted residuals.

To adjust for covariates in a general way, we propose to perform a kernel principal component analysis (kernel PCA) [25], a general extension of regular PCA, on the constructed phenotype kernel matrix and treat the resulting kernel PCs as surrogate phenotypes. We can then regress out the covariates from the kernel PCs and reconstruct the phenotype kernel matrix with the adjusted PCs. The same procedure is performed on the genotype kernel matrix. After algebraic manipulation (see Derivation of covariate-adjusted KRV coefficient), the adjusted KRV coefficient is of the form:

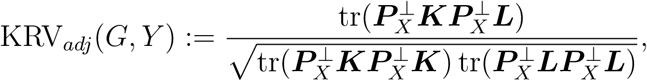

where 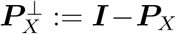 and ***P***_*X*_ is the projection matrix onto the column space of ***X***. We adjust for covariates on both the phenotype kernel and the genotype kernel, due to the symmetry of the KRV coefficient. Similar to the analogy between KRV and the sample correlation, we can view the adjusted KRV coefficient as an extension of the sample partial correlation [26, 27]. The usual hypothesis testing procedure in the KRV framework can be applied to the adjusted KRV statistic to obtain a p-value. In this case, the null hypothesis is that there is no association between the genetics and the phenotypes after adjusting for the effects of the covariates.

### Application of covariate-adjusted KRV to HCHS/SOL

To identify genetic variants associated with the overall gut microbiome composition in Hispanic/Latino individuals, we applied the covariate-adjusted KRV test to the HCHS/SOL study. HCHS/SOL is a community-based cohort study aimed to identify factors that affect the health of Hispanic/Latino individuals. The study recruited 16,415 Hispanic/Latino adults of diverse ethnic background from four U.S. metropolitan areas (Bronx, NY; Chicago, IL; Miami, FL; San Diego, CA) [20]. Genome-wide DNA sequencing data were available in 12,803 participants. As an ancillary study, the HCHS/SOL Gut Origins of Latino Diabetes (GOLD) study was further conducted to investigate the role of gut microbiome composition in health outcomes such as diabetes in Hispanic/Latino individuals [21], where gut microbiome profiles were available in 1674 participants (a subgroup of the HCHS/SOL participants) based on 16S rRNA gene sequencing. More details on collection and pre-processing of genetic and microbiome data in HCHS/SOL are provided in Description of the HCHS/SOL study.

We considered genetic variants (including both single-nucleotide polymorphisms, or SNPs, and insertion/deletion variants, or indels) within *±*10 kb of gene regions and grouped the variants into gene-level variant-sets correspondingly. The microbiome operational taxonomic units (OTUs) were collapsed at the genus level and rarefied to accommodate differential read depth. We used a linear kernel for the genetic data and different kernels for the microbiome data, including Bray-Curtis, unweighted UniFrac, weighted UniFrac and generalized UniFrac (see Choice of kernels for details on these kernels). For each gene, we assessed the association between the common variants (with minor allele frequency, or MAF, *≥* 0.05) within the gene and the community-level microbiome profile, using both adjusted and unadjusted KRV tests. In the adjusted KRV, we mainly controlled for the top 5 PCs of genome-wide genetic variability (denoted as the PC-adjusted KRV), as they well captured the population structure of the sample. Individuals from different populations and ethnic groups often have systematic differences in their genetic and microbiome profiles [28, 29], so population structure is an important confounder in our analysis. We also performed additional analyses that adjusted for other non-confounding covariates including age, gender and study sites.

Our investigation of the genetic effect on the microbiome involved two stages. In the first stage, we tested the association between the variants in each gene and the microbiome profile at the community level. In the second stage, for any genes called significant in the first stage, we marginally assessed the association between each of the individual variants within those genes and the community-level microbiome profile to look for significant variants, using the covariate-adjusted KRV. Bonferroni correction was applied in both stages. Since this was a nested hypothesis testing approach, the second-stage test only required correction for the number of variants in the genes that were called significant in the first stage. All analyses were performed on unrelated individuals (pairwise kinship coefficient *≤* 0.05) where genetic data, microbiome data and covariates data were available.

As a result, we performed our analyses on 1219 unrelated participants from HCHS/SOL where all relevant data were available. Among these individuals, 47.0% identified their background as Mexican, 14.8% as Cuban, 12.7% as Puerto Rican, 10.3% as Central American, 7.7% as South American and 7.5% as Dominican. Microbiome count data were obtained on 408 genera, rarefied to 10,000 total counts per individual. A total of 19223 gene-level variant-sets that contained at least one common variant were available. Figure 1 shows the p-value QQ-plots of the first-stage (gene-level) analysis results. For all microbiome kernels, the unadjusted KRV produces highly anti-conservative p-values (with large genomic inflation factors), while the PC-adjusted KRV has well-controlled type I error rates (with genomic inflation factors *≤* 1.05), confirming that population structure is the major confounder in our study. The gene-level Manhattan plots based on the PC-adjusted KRV are shown in Supplementary Figure S1.

**Figure 1:**
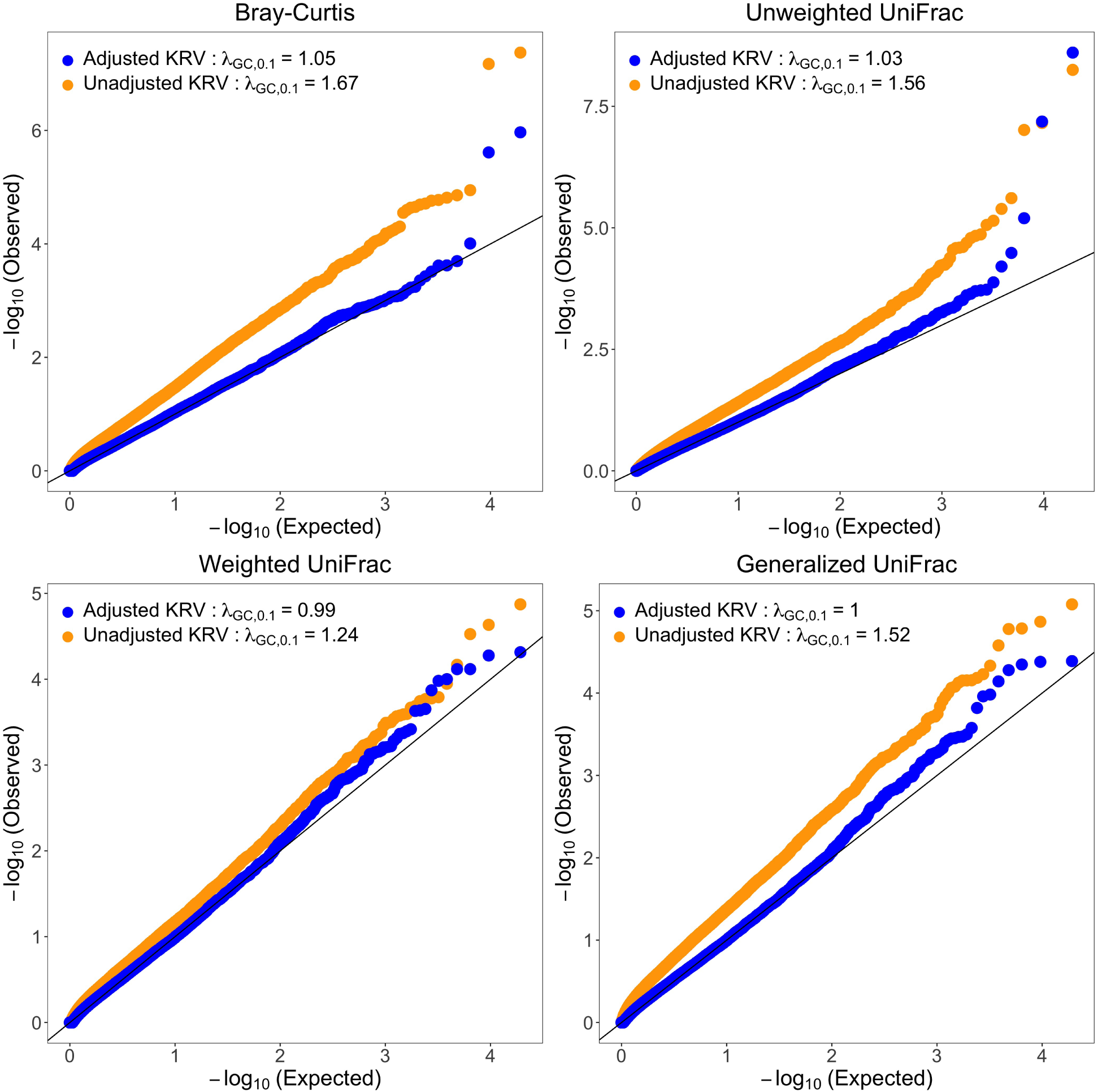
P-value QQ-plots from the first-stage (gene-level) analysis of the HCHS/SOL data. Each QQ-plot corresponds to a distinct microbiome kernel. In the adjusted KRV, the top 5 PCs of genome-wide genetic variability were adjusted. *λ*_*GC*,0.1_ represents the genomic inflation factor evaluated at the 10th percentile.

Table 1 shows the genes identified at a genome-wide significance in the PC-adjusted first-stage analysis (*α* = 0.05*/*19223 = 2.6 *×* 10^*−*6^). We have found two genes, *IL23R* and *C1orf141*, using the Bray-Curtis kernel and two genes, *MTMR12* and *ZFR*, using the unweighted UniFrac kernel. When the analysis is performed on a reduced set of individuals (*n*=1096) where additional covariates (age, gender and study sites) are available and adjusted, *IL23R* and *C1orf141* are no longer genome-widely significant (Supplementary Table S1); similar results are observed for a PC-adjusted analysis on the same subsample. To investigate the reason for this power loss, we perform PC-adjusted analyses on random sub-samples of the same size from the original 1219 individuals. Around half of the times, at least two out of the four genes no longer have genome-wide significance, indicating that the non-significant results in the reduced sample are likely due to sample size loss. Nevertheless, the results from the two adjusted analyses are qualitatively similar.

**Table 1:** Significant genes identified from the first-stage (gene-level) analysis of the HCHS/SOL data, using the PC-adjusted KRV (*α* = 2.6 *×* 10^*−*6^).

| Microbiome kernel | Significant genes | Number of common variants | P-value |
| --- | --- | --- | --- |
| Bray-Curtis | *C1orf141* | 484 | $1.1 \times 10^{-6}$ |
| | *IL23R* | 284 | $2.4 \times 10^{-6}$ |
| Unweighted UniFrac | *MTMR12* | 174 | $6.5 \times 10^{-8}$ |
| | *ZFR* | 288 | $2.5 \times 10^{-9}$ |

The top 5 PCs of genome-wide genetic variability were adjusted.

Among these genes, *IL23R* is of considerable interest: it encodes one part of the receptor for interleukin-23 (IL-23), a pro-inflammatory cytokine closely involved in autoimmunity [30]. The *IL23R* gene has been associated with inflammatory bowel diseases (IBD) including Crohn’s disease and ulcerative colitis [31, 32]. In a previous genetic association study of microbiome composition [33], the protective variant of the *IL23R* gene (rs11209026) was associated with a higher microbiome diversity and richness and a higher abundance of beneficial gut bacteria in the ileum of healthy individuals, suggesting the influence of host genetics on the microbiome prior to onset of IBD. In addition, a mouse-based experimental study [34] showed that mice deficient in intestinal *IL23R* expression had altered gut microbiota and were susceptible to colonic inflammation, where increased disturbance of gut microbiota exacerbated the disease activity. Coupled with these results, our finding further supports that the gut microbiome may mediate the host genetic effect on the development of inflammatory diseases like IBD. In its normal function, the *IL23R* gene likely helps shape the overall gut microbiota towards a healthy composition, which may in turn support normal immune activities and prevent gut inflammation.

The other genes are also interesting to further explore. The *C1orf141* gene, with uncharacterized protein function, has overlapping regions with *IL23R*. Variants in the *IL23R-C1orf141* region have been associated with susceptibility to Vogt-Koyanagi-Harada disease, a multi-system autoimmune disorder that affects pigmented tissues, in Chinese and Japanese populations [35, 36]. The *ZFR* gene encodes the highly conserved zinc finger RNA-binding protein, which is shown to prevent excessive type I interferon activation by regulating alternative pre-mRNA splicing [37]. Prevention of excessive type I interferon activation is important for the regulation of immune responses. The *MTMR12* gene encodes an adapter protein for myotubularin-related phosphatases and is likely involved in skeletal muscle functions [38]. Overall, most of the significant genes have a role in immunity, indicating an interaction between the host genetics and the gut microbiome in facilitating immune responses or developing autoimmune disorders.

Figure 2 shows the Manhattan plots and linkage disequilibrium (LD) heatmaps from the second-stage analysis of the HCHS/SOL data, using the PC-adjusted KRV. The *IL23R* and *C1orf141* genes were combined into a single *IL23R-C1orf141* region due to overlapping variants. Based on the analysis using the Bray-Curtis kernel, there are 72 significant variants (out of 557 common variants) in the *IL23R-C1orf141* region (*α* = 0.05*/*557 = 8.98 *×* 10^*−*5^). Based on the analysis using the unweighted UniFrac kernel, there are 114 significant variants (out of 288 common variants) in *ZFR* and 125 significant variants (out of 174 common variants) in *MTMR12* (*α* = 0.05*/*(288 + 174) = 1.08 × 10^*−*4^). Relevant information including positions, rsID and p-values for these variants is reported in Supplementary Table S2. From the LD heatmaps, in each gene, the significant variants share a high level of linkage disequilibrium with one other. Future fine mapping of causal variants that affect the microbiome composition will be needed.

**Figure 2:**
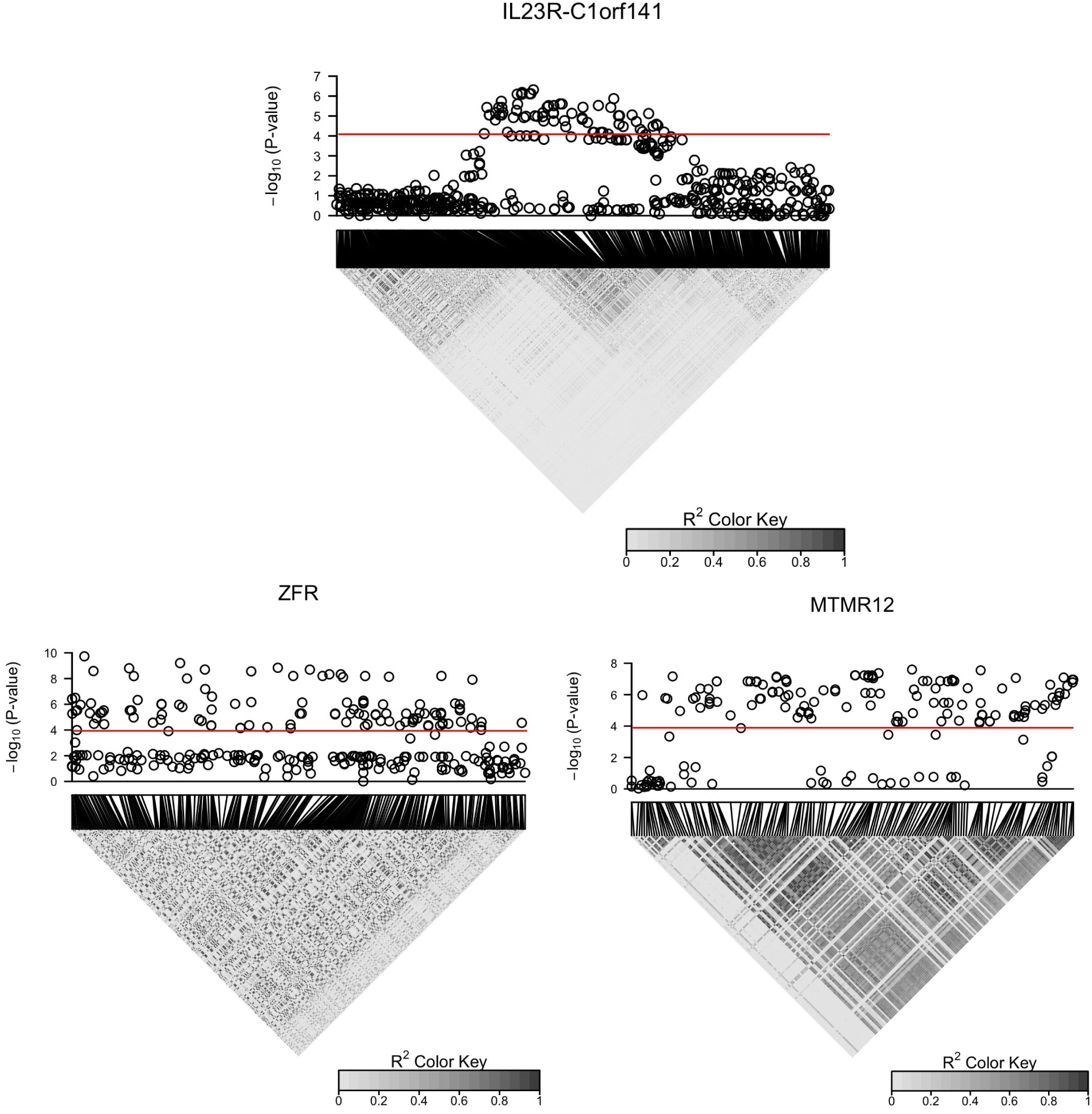
Manhattan plots and linkage disequilibrium (LD; *R*^2^) heatmaps from the second-stage (variant-level) analysis of the HCHS/SOL data, using the PC-adjusted KRV. The Bray-Curtis kernel was used for analysis of variants in the *IL23R-C1orf141* region; the unweighted UniFrac kernel was used for analysis of variants in *ZFR* and *MTMR12*. The top 5 PCs of genome-wide genetic variability were adjusted. The red lines represent variant-level signficance after Bonferroni correction (*α* = 8.98 ×10^*−*5^ for variants in the *IL23R-C1orf141* region, and 1.08 × 10^*−*4^ for variants in *ZFR* and *MTMR12*). A large *R*^2^ value indicates high LD.

To confirm the validity of the adjusted KRV approach, we further conduct kernel PCA on the Bray-Curtis and unweighted UniFrac kernel matrices, and check whether individuals’ microbiome profiles, captured by the top two kernel PCs, differ by genotypes of the top (most significant) variant from each identified gene. This is similar to a PCoA analysis. Figure 3 shows that, for each top variant, the 95% confidence ellipses for different genotypes are well separated from one other, corroborating the findings by the adjusted KRV.

**Figure 3:**
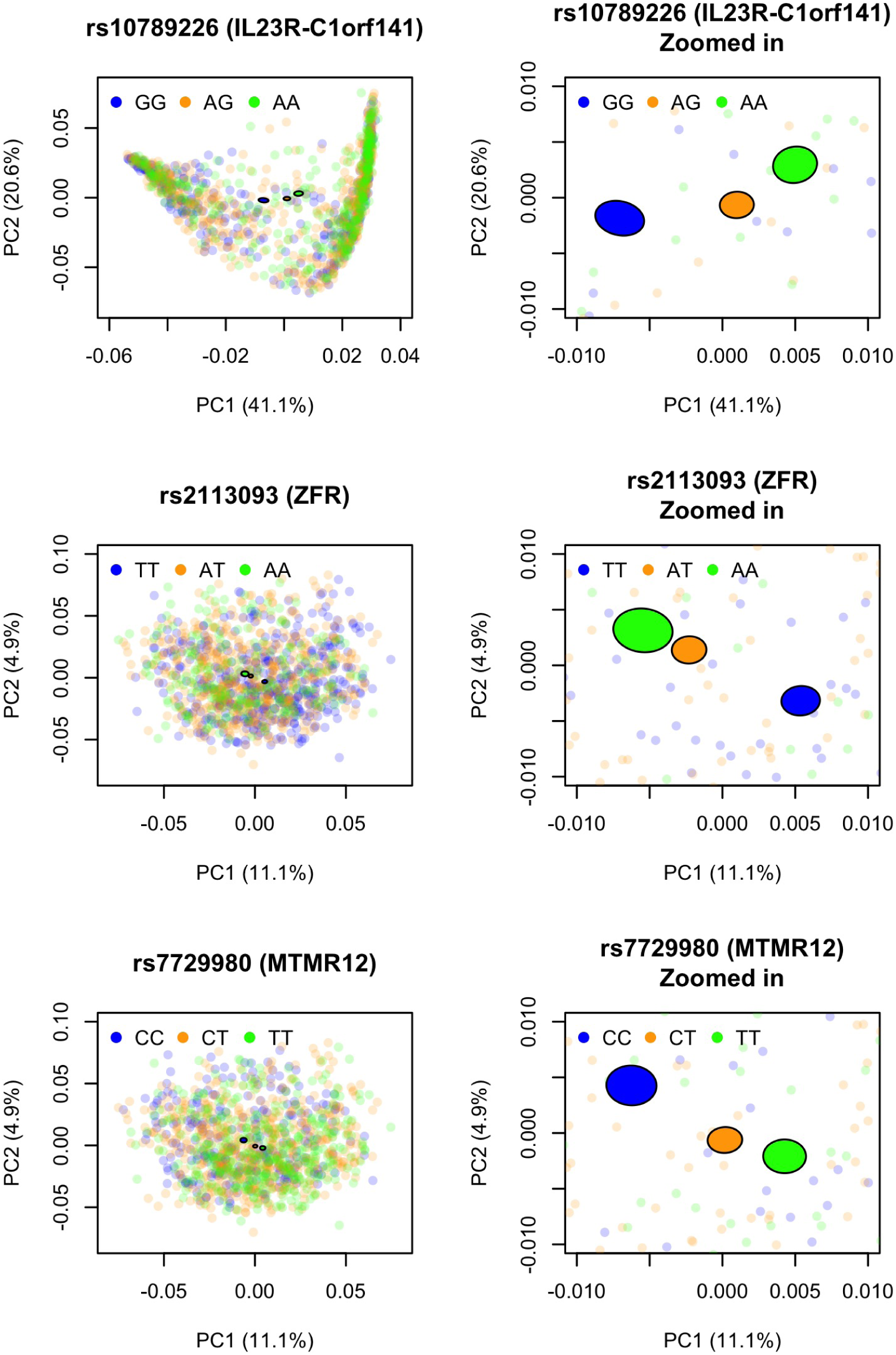
PC2 vs. PC1 from kernel PCA on the microbiome kernel, colored by the genotype of top variants from the significant genes in the HCHS/SOL study. For each variant, a 95% confidence ellipse (shown as a filled ellipse with black borders) was constructed for individuals from each genotype. The Bray-Curtis kernel was used for the top variant in the *IL23R-C1orf141* region; the unweighted UniFrac kernel was used for the top variants in *ZFR* and *MTMR12*. The percent of variance captured by each PC was provided in the axis labels.

We next examine the replication of signals found by previous GWAS studies in our analysis. Kurilshikov et al. [5] analyzed a sample of 18,340 individuals that comprised of 24 multi-ancestry cohorts, including the HCHS/SOL GOLD cohort. They reported an association between the *LCT* locus (rs182549) and *Bifidobacterium* abundance at a study-wide significance (p-value = 1.28 × 10^*−*20^). In our gene-level analysis using the PC-adjusted KRV, the *LCT* gene is nominally significant based on the unweighted UniFrac kernel (p-value = 0.013), but not significant at the genome-wide level. In addition, we have examined the significance of 63 previously reported genes that harbor variants associated with microbiome beta-diversity [10, 12, 14, 39, 40] (Supplementary Table S3). 59 out of 63 genes include at least one common variant in the HCHS/SOL data. Two genes are replicated with nominal significance: *BANK1* based on the unweighted UniFrac kernel (p-value = 0.017) and the weighted UniFrac kernel (p-value = 0.046), and *MAST3* based on the weighted UniFrac kernel (p-value = 0.041) and the generalized UniFrac kernel (p-value = 0.049). *BANK1* is associated with systemic lupus erythematosus and *MAST3* is associated with IBD, corroborating the role of immunity-related genes in shaping gut microbiota. However, none of the genes are significant at the genome-wide level.

### Simulation studies

We conducted simulation studies to further evaluate the type I error rate and power of the covariate-adjusted KRV test. We simulated genotype data and microbiome OTU count data under realistic settings, and introduced population stratification as a confounder that affected both genetic and microbiome data. Detailed simulation procedures are provided in Methods: Simulation studies.

The general simulation setting is as following. We considered a sample size of 1000. SNP genotype data were simulated over a 1 Mb chromosome region for 500 individuals of African ancestry and 500 individuals of European ancestry. Count data of 856 microbiome OTUs were simulated using a Dirichlet-multinomial distribution. To introduce population structure into the OTU count data, we increased the relative abundance of the 10 most common OTUs by 10% in African individuals. Both unadjusted and adjusted KRV tests were performed to test the association between the overall microbiome composition and common SNPs (with MAF *≥* 0.05) within an 8 kb subregion of the 1 Mb chromosome. In the adjusted KRV test, the top PC of genetic variability (obtained from PCA on SNP data over the entire 1 Mb region) was used as the covariate, a surrogate for population structure. We used a linear kernel for genetic data and four different kernels for microbiome data: Bray-Curtis, unweighted UniFrac, weighted UniFrac and generalized UniFrac.

To evaluate the type I error rate, we used the above simulation setting without introducing any genetic effect on the microbiome; 10,000 data sets were simulated. To evaluate the statistical power of the adjusted KRV, we introduced genetic effect on the microbiome in three different scenarios, on top of the general simulation setting. In all three scenarios, we simulated a pleiotropy effect, where a single SNP affected the abundance of multiple microbiome OTUs. In Scenario 1, a single SNP affected the abundance of the 11th - 20th most common OTUs. In Scenario 2, a single SNP affected the abundance of OTUs from a relatively common phylogenetic cluster. In Scenario 3, a single SNP affected the abundance of 5 rare OTUs. For all scenarios, we considered both small and large effect sizes. We also performed two competing methods based on univariate microbiome phenotypes: linear regression and SNP-set kernel association test (SKAT) [17] (see Methods: Simulation studies for details). For each power scenario, 1000 data sets were simulated.

Table 2 shows the empirical type I error rates of both unadjusted and adjusted KRV tests at different significance levels. The unadjusted KRV has inflated type I error rates for all microbiome kernels except unweighted UniFrac. In contrast, the adjusted KRV maintains valid type I error rates for all microbiome kernels. Note that in our simulation setting, population structure affected the abundance of common OTUs, which was unlikely to change these OTUs’ presence. Since the unweighted UniFrac kernel only captures presence/absence, but not abundance information of a taxon, the population stratification of microbiome profiles is not reflected in the unweighted UniFrac kernel. This absence of confounding effect leads to a valid type I error rate for the unweighted UniFrac kernel even when the unadjusted KRV is used.

**Table 2:** Empirical type I error rate of unadjusted and covariate-adjusted KRV at nominal level *α* under simulation.

| Method | Microbiome kernel | $\alpha$ | | |
| --- | --- | --- | --- | --- |
|  |  | 0.05 | 0.01 | 0.001 |
| Unadjusted KRV | Bray-Curtis | 0.2403 | 0.0936 | 0.0255 |
|  | Unweighted UniFrac | 0.0484 | 0.0094 | 0.0011 |
|  | Weighted UniFrac | 0.1371 | 0.0371 | 0.0057 |
|  | Generalized UniFrac | 0.1412 | 0.0416 | 0.0063 |
| Adjusted KRV | Bray-Curtis | 0.0473 | 0.0114 | 0.0012 |
|  | Unweighted UniFrac | 0.0523 | 0.0115 | 0.0009 |
|  | Weighted UniFrac | 0.0507 | 0.0095 | 0.0012 |
|  | Generalized UniFrac | 0.0499 | 0.0097 | 0.0011 |

Linear kernel was used for genetic data.

Figure 4 shows the empirical power of the covariate-adjusted KRV test and competing methods under small effect sizes, at the nominal level *α* = 0.05. In general, for each power scenario, the adjusted KRV has a much higher power than linear regression and SKAT, regardless of the microbiome kernel being used (with the exception of unweighted UniFrac in Scenario 1 and 2). Next we focus on the adjusted KRV and compare across microbiome kernels: in Scenario 1, the Bray-Curtis kernel has the highest power; in Scenario 2, the weighted UniFrac kernel has the highest power; in Scenario 3, the unweighted UniFrac kernel has the highest power. These results are consistent with the ways these microbiome similarity measures are constructed. The Bray-Curtis kernel is efficient in detecting abundance changes in common OTUs. The weighted UniFrac kernel has more power to detect abundance changes in common phylogenetic clusters, and the unweighted UniFrac kernel is more efficient in detecting changes in rare lineages. Again, due to the nature of unweighted UniFrac, all three methods based on this kernel have little power in Scenario 1 and 2, where the SNP effect on common OTUs or common phylogenetic clusters is unlikely to change their presence.

**Figure 4:**
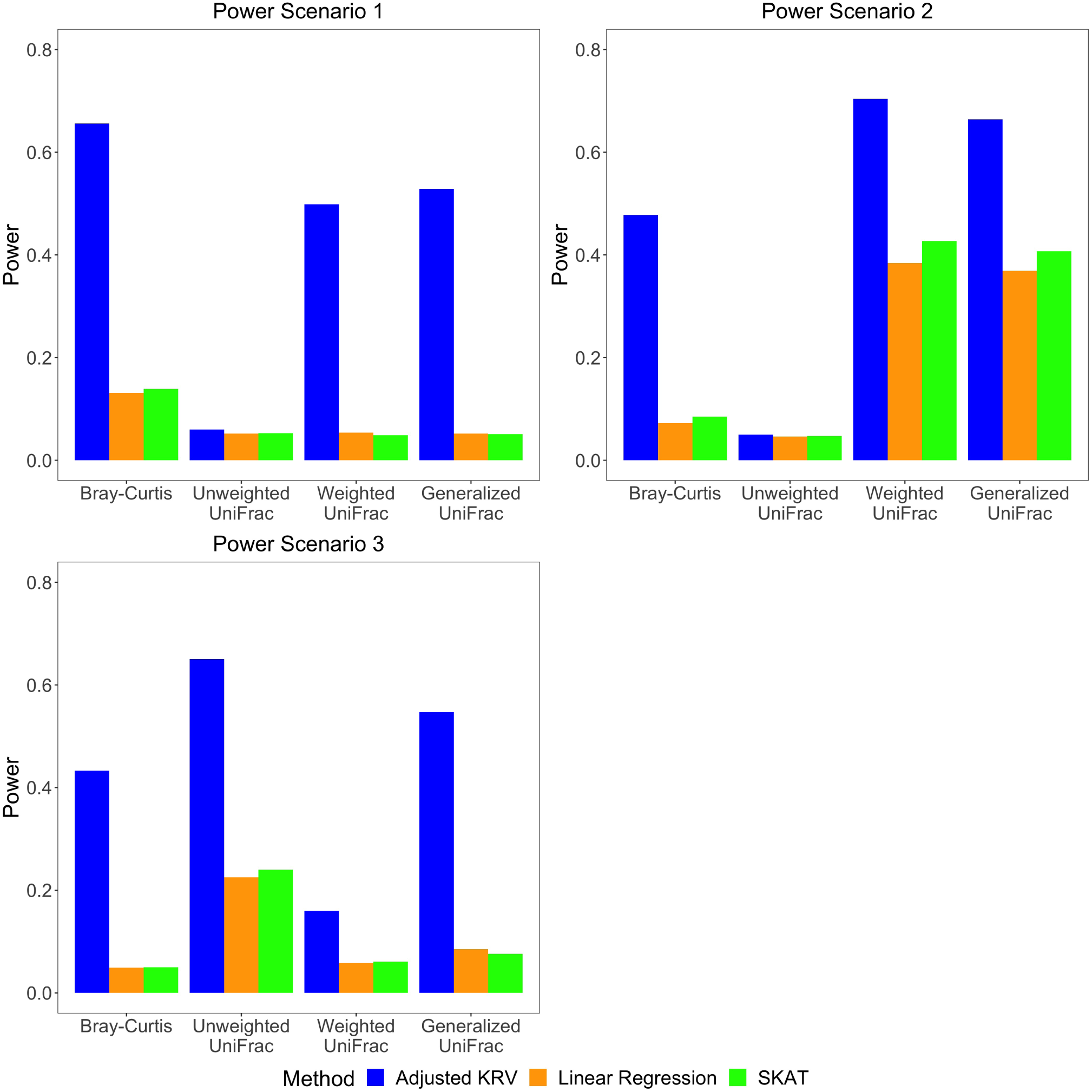
Empirical power of covariate-adjusted KRV and competing methods at nominal level *α* = 0.05 for different microbiome kernels under small effect sizes. Linear kernel was used for genetic data.

Under large effect sizes (Supplementary Figure S2), while the covariate-adjusted KRV displays a clear improvement in power, the overall patterns are similar to those under small effect sizes.

## Discussion

We have introduced the covariate-adjusted KRV, a novel microbiome GWAS approach to evaluate the association between a group of genetic variants at the gene level and the overall microbiome composition at the community level, while adjusting for covariates. Simulation studies show that the covariate-adjusted KRV maintains valid type I error rates in the presence of confounders and has a much higher power compared to other microbiome GWAS methods that rely on univariate microbiome phenotypes. In a genome-wide analysis of the HCHS/SOL data, we have identified four genes associated with microbiome beta-diversity. We have also identified specific variants within these genes in a second-stage analysis, which will be useful for future ascertainment of causal variants that affect the gut microbiota.

Most of the identified genes based on the HCHS/SOL data have been previously implicated in immune functions or immunity-related disorders. This is consistent with the works by Blekhman et al. [6] and Rühlemann et al. [12], where loci in immunity-related genes and pathways have been shown to correlate with gut microbiome composition. The *IL23R* gene is especially interesting for future study, due to its recognition in previous microbiome genetic association studies [33] and its role in IBD, a chronic inflammatory disease that involves both genetic and microbial factors. Many genetic markers associated with IBD are involved in the interactions between the immune system and the microbiome [41, 42]. Furthermore, IBD is characterized by shift in the gut microbiome composition [43, 44], and specific microbes have also been shown to predict response to therapy [45] and postoperative disease recurrence [46] in patients with IBD. Therefore, our finding supports previous work and could contribute to future investigation of the disease etiology. Finally, as HCHS/SOL is one of the most comprehensive studies of Hispanic/Latino populations in the US, the results from our analysis will help inform important genetic risk factors for gut-microbiome-related health outcomes in Hispanic/Latino individuals.

Although the covariate-adjusted KRV has valid type I error rates regardless of the kernels used, selecting appropriate kernels that reflect the actual patterns of association is important for maintaining a good statistical power. Different kernels measure different aspects of the structure within the data and assume different association patterns. In the analysis of the HCHS/SOL data, using different microbiome kernels, we discovered distinct significant genes. This is likely because these genes affect different aspects of the microbiome composition. For example, loci in the *IL23R-C1orf141* region, identified using Bray-Curtis, likely affect abundances of common microbial taxa such as *Bacteroides* and *Prevotella* [21]. Loci in *ZFR* and *MTMR12*, identified using unweighted UniFrac, likely affect the presence/absence of certain rare microbial lineages. Often we do not have prior knowledge on the ways genetics is associated with the microbiome. A possible extension would be to use an omnibus test that accommodates multiple possible kernels. For example, as proposed by Zhan et al. [19], we could construct an omnibus kernel matrix via a weighted sum of multiple candidate kernel matrices. Another approach would be to combine p-values obtained using different candidate kernels into a single p-value, such as the Cauchy p-value combination method [47].

We have also investigated the replication of signals from previous microbiome GWAS studies. The multi-cohort sample used by Kurilshikov et al. [5] includes the HCHS/SOL GOLD cohort. While Kurilshikov et al. reported an association between the *LCT* locus (rs182549) and *Bifidobacterium* abundance at a study-wide significance, the *LCT* gene was not identified as genome-widely significant in our analysis. *Bifidobacterium* was a relatively common genus (representing 1.04% abundance of all microbial genera) in the HCHS/SOL data. However, when we used microbiome kernels that are efficient in detecting abundance changes in common taxa, such as Bray-Curtis and weighted UniFrac, abundance differences in *Bifidobacterium* were likely overshadowed by those in the most common genera such as *Bacteroides* and *Prevotella* (representing 23.7% and 25.0% abundances of all microbial genera, respectively). This discrepancy in results might reflect the difference between taxon-level and community-level analyses.

Two previously reported beta-diversity-associated genes [10] have been replicated in our analyses at a nominal significance, but none of the previous signals [10, 12, 14, 39, 40] reaches genome-wide significance. There are several possible reasons. First, compared to environmental effect, most host genetic influences on microbiome composition have relatively small effect sizes [3]. The sample sizes of current microbiome GWAS studies, including our study, are still too small to achieve enough statistical power. Second, there is considerable variation across studies in the collection and processing of microbiome data, leading to difficulties in reproducibility. Lastly, certain genetics-microbiome associations might be specific to ancestry or populations. In addition, since we focused on genetic loci within or close to gene regions, we were unable to evaluate the significance of previously identified loci that fell in intergenic regions.

In conclusion, we have proposed a promising approach to study the covariate-adjusted association between host genetic variation and community-level microbiome composition, which demonstrates good performances in both simulations and real data analysis. The genes and loci identified using our approach will help elucidate the complex interactions among host genetics, gut microbiome and host immune systems. With the increasing occurrences of high-dimensional traits in large-scale genetic association studies, we expect the covariate-adjusted KRV to bring more discoveries by taking advantage of the innate structure within the genetic and phenotypic data.

## Methods

### Choice of kernels

In the KRV framework, kernel functions are used to summarize pairwise similarities in genotype and phenotype profiles among the subjects. In order to improve the statistical power in hypothesis testing, we would like to choose kernels that better reflect the actual structure within the genetic and phenotype data as well as the patterns of association [15, 48]. Theoretically, for the KRV statistic in (1) to be well-defined, the kernel matrices need to be positive semi-definite. We now review some of the common kernels used for genetic and microbiome data, respectively.

For genotype data, popular kernel functions include the linear kernel 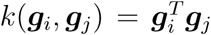 and the identity-by-state (IBS) kernel 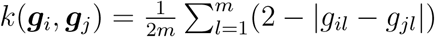. The linear kernel assumes that the genetic variants are associated with the traits in a linear fashion. The IBS kernel defines pairwise similarity as the pairwise genotype matching averaged over all genetic variants, and is useful when there are epistatic effects among the variants [17]. Depending on analysis interests (e.g. rare-variant analysis), it is also possible to incorporate a weight for each variant in the linear and IBS kernels [17].

For microbiome data at the community level, the kernel matrix can be obtained by transforming known ecological or phylogenetic dissimilarity measures (i.e., beta-diversity measures). For example, Bray-Curtis dissimilarity quantifies the dissimilarity between two microbial communities based on the difference in counts at each taxon between the two communities. The UniFrac distances are dissimilarity measures based on the phylogenetic structure of the taxa [49, 50, 51]: the unweighted UniFrac distance is calculated as the fraction of branch lengths within the phylogenetic tree that are not shared between the two communities; the weighted UniFrac distance further incorporates taxa abundance information on the basis of the unweighted distance; the generalized UniFrac distance is a compromise between weighted and unweighted UniFrac distances. For a given pairwise dissimilarity matrix ***D***, the corresponding kernel matrix can be constructed as:

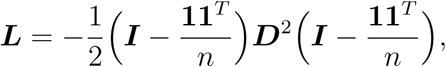

where ***D***^2^ is the element-wise square of ***D***. To ensure that the kernel matrix ***L*** is positive semi-definite, we further apply a correction procedure as implemented in the MiRKAT R package [15], where we perform an eigendecomposition of ***L***, convert any negative eigenvalues to zero and then reconstruct the kernel matrix.

### Derivation of covariate-adjusted KRV coefficient

Suppose that we have a phenotype kernel matrix ***L*** and a full-rank covariates matrix ***X*** that includes a column of 1’s. We first perform a kernel PCA (equivalent to an eigendecomposition) on the phenotype kernel matrix and obtain a matrix **Φ** such that:

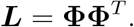

Here each column of **Φ** is a kernel principal component (kernel PC) of ***L*** and has the form 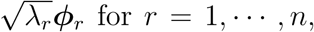, where *λ*_*r*_ is the *r*th eigenvalue of ***L*** and ***ϕ***_*r*_ is the corresponding eigenvector for *λ*_*r*_. We can view **Φ** as a finite sample basis for the space spanned by the phenotype kernel function *𝓁* (*·,·*).

We then regress out the covariates ***X*** from each kernel PC:

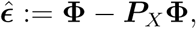

where ***P***_*X*_ = ***X***(***X***^*T*^ ***X***)^*−*1^***X***^*T*^ is the projection matrix onto the column space of ***X***. Now 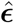 represents a sample basis that is orthogonal to the covariates ***X***. We can construct a new phenotype kernel matrix from this residual basis: 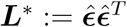. Note that ***L***^*∗*^ can be expressed in terms of ***L***:

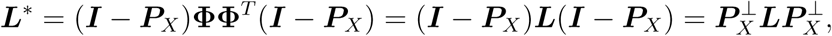

where we let 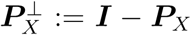. Similar procedures can be performed on the genotype kernel matrix ***K*** to obtain the adjusted genotype kernel matrix 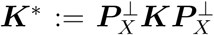. Both ***K***^*∗*^ and ***L***^*∗*^ are column-centered, since the covariates matrix ***X*** includes a column of 1’s, accounting for the intercept in a regression. We can then construct a KRV statistic from the adjusted kernel matrices ***K***^*∗*^ and ***L***^*∗*^:

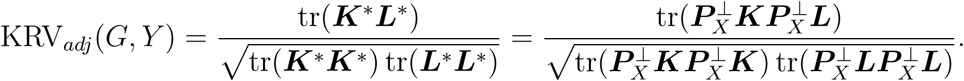

Such a strategy of covariate adjustment can be seen as a special case of conditional independence (or uncorrelatedness) testing in a kernel-based framework, as proposed by Zhang et al. and Strobl et al. [52, 53]. In the context of microbiome GWAS, we are testing the correlation between genetic variants and microbiome community profiles, while conditioning on the covariates.

### Description of the HCHS/SOL study

HCHS/SOL is a community-based prospective cohort study aimed to identify risk factors for health outcomes in Hispanic/Latino individuals. The study recruited 16,415 Hispanic/Latino adults aged 18 - 74 years at four U.S. field centers (Bronx, NY, Chicago, IL, Miami, FL, and San Diego, CA), using a two-stage probability sampling design [20].

12,803 participants consented to genetic studies. Genotyping was performed on an Illumina custom array, SOL HCHS Custom 15041502 B3, which consisted of the Illumina Omni 2.5M array (HumanOmni2.5-8v1-1) and *∼*150,000 custom SNPs [54]. Quality control, genotype imputation and estimation of pairwise kinship coefficients and PCs of genome-wide genetic variability were described in detail by Conomos et al. [54]. In addition to the quality control procedures described in [54], prior to the microbiome GWAS analysis, we also filtered imputed genetic variants based on an “effective minor allele count”: 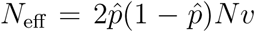, where 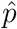 is the estimated minor allele frequency, *N* is the sample size and *v* is the ratio of observed variance of imputed dosages to the expected binomial variance [55]. We retained variants with sufficient minor allele counts and excluded any variants with *N*_eff_ *<* 30.

Gut microbiome profiles were available in 1674 participants, a subset of the HCHS/SOL participants, from the HCHS/SOL GOLD ancillary study. Based on the collected stool samples, DNA extraction and 16S rRNA gene sequencing were performed according to the Earth Microbiome Project (EMP) standard protocols [56]. Subsequent bioinformatic processing of the microbiome sequencing data was described in detail by Kaplan et al. [21].

The HCHS/SOL study was approved by the Institutional Review Boards of all participating institutions, and written informed consent was obtained from all participants.

### Simulation studies

To simulate genotype data with population structure, we first generated 10,000 haplotypes of African ancestry and another 10,000 haplotypes of European ancestry over a 1 Mb chromosome according to coalescent theory using the *cosi2* program [57]. To form a sample, we then generated the genotype of each African individual in the sample by randomly selecting and pairing 2 haplotypes from the 10,000 founding African haplotypes. A similar procedure was used to generate the genotypes of European individuals.

We used a Dirichlet-multinomial distribution to generate microbiome OTU counts for each individual in the sample, as this distribution well accommodates the over-dispersion of microbiome count data [15, 58]. To ensure a realistic simulation of OTU counts, we estimated the parameters of the Dirichlet-multinomial distribution from a real upper-respiratory-tract microbiome data set [59], which consisted of 856 OTUs. This data set is publicly available as part of the GUniFrac R package. We assumed 1000 total OTU counts per individual. After introducing population structure into the OTU count data by increasing the counts of the 10 most common OTUs by 10% in African individuals, we rarefied the OTU counts back to 1000 total counts per individual. Here we used the estimated mean proportion parameters of the Dirichlet-multinomial distribution as a measure of OTU prevalence.

To evaluate the power of the covariate-adjusted KRV, we introduced an association between the genetics and the microbiome in three difference scenarios. Let *g*_*i*_ be the genotype (0, 1 or 2) of individual *i* at a chosen common SNP (with MAF *≥* 0.05). In Scenario 1, for each individual *i*, we increased the counts of the 11th - 20th most common OTUs by a factor of *f*_*i*_, where *f*_*i*_ = 1 + *c*_1_*g*_*i*_. In Scenario 2, utilizing the available phylogenetic tree for the 856 OTUs [59], we increased the counts of OTUs from a relatively abundant cluster (representing 10.3% abundance of the total OTU counts) by a factor of *f*_*i*_ for each individual *i*, where *f*_*i*_ = 1 + *c*_2_*g*_*i*_. In Scenario 3, for each individual *i*, we increased the counts of 5 rare OTUs (chosen randomly from the top 40 rarest OTUs) by an addition of *a*_*i*_, where *a*_*i*_ = *c*_3_*g*_*i*_. We considered two sets of effect sizes: (a) small effect sizes: *c*_1_ = *c*_2_ = 0.3, *c*_3_ = 0.5 and (b) large effect sizes: *c*_1_ = 0.8, *c*_2_ = 0.7, *c*_3_ = 1. After introducing these genetic effects on the microbiome, we again rarefied the OTU counts to 1000 total counts per individual.

In the power simulation, we considered two competing methods that rely on univariate microbiome phenotypes. The first method was linear regression, where we performed kernel PCA on both genotype and microbiome kernel matrices and regressed the top PC of the microbiome kernel on the top PC of the genotype kernel, while adjusting for covariates. The second method was SNP-set kernel association test (SKAT) [17], a kernel machine regression framework for assessing the general association between a univariate trait and multiple genetic variants. Here we performed kernel PCA on the microbiome kernel matrix and used the SKAT test to regress the top PC of the microbiome kernel on the genetic variants within the pre-specified region, while adjusting for covariates; a linear kernel was used for genetic data in the SKAT test.

### Computation time

We estimated the computation time of the covariate-adjusted KRV test for different sample sizes. For each sample size, we simulated 10 data sets and reported the average computation time. Given constructed genotype and microbiome kernel matrices and 10 covariates, the average computation times are 0.09, 1.23, 12.58 and 97.57 seconds on a laptop (2.7 GHz CPU and 16 GB memory) for sample sizes of 200, 500, 1000 and 2000, respectively. The gene-level analysis of the HCHS/SOL data set (with one genotype kernel, 4 microbiome kernels and 19223 variant-sets) took approximately 6 hours on a high-performance computing cluster (each node with 24 cores, 3.00 GHz CPU and 384 GB memory), with computing jobs divided by chromosome.

### Web resources

Figure 2 was produced using the LDheatmap R package v1.0: https://cran.r-project.org/web/packages/LDheatmap. The 95% confidence ellipses in Figure 3 were produced using the ordiellipse() function of the vegan R package v2.5: https://cran.r-project.org/web/packages/vegan. The covariate-adjusted KRV test is implemented as part of the KRV() function in the MiRKAT R package v1.2.1: https://cran.r-project.org/web/packages/MiRKAT. Other tools include: *cosi2* program: https://software.broadinstitute.org/mpg/cosi2. SKAT R package v2.0.1: https://cran.r-project.org/web/packages/SKAT. GUniFrac R package v1.2: https://cran.r-project.org/web/packages/GUniFrac.

## Data availability

The HCHS/SOL data used in our study are deposited at the database of Genotypes and Phenotypes (dbGap; http://view.ncbi.nlm.nih.gov/dbgap) and Biologic Specimen and Data Repository Information Coordinating Center (BIOLINCC; https://biolincc.nhlbi.nih.gov). The genotype and covariates data are available at dbGap under accession codes: phs000880.v1.p1 and phs000810.v1.p1. The 16S rRNA gene sequences are deposited in QIITA (https://qiita.ucsd.edu) under ID 11666, and European Nucleotide Archive (ENA; https://www.ebi.ac.uk/ena) under accession code ERP117287. HCHS/SOL has established a procedure for the scientific community to apply for access to participant data, with such requests reviewed by the Steering Committee of the HCHS/SOL project. These policies are described at https://sites.cscc.unc.edu/hchs.

## Code availability

The covariate-adjusted KRV approach is implemented as part of the KRV() function in the MiRKAT R package v1.2.1, available at the Comprehensive R Archive Network (CRAN): https://cran.r-project.org/web/packages/MiRKAT. Instructions for usage and codes for reproduction of simulation results in this study are available at https://github.com/pearl-liu/Covariate-Adjusted-KRV.

## Acknowledgements

The Hispanic Community Health Study/Study of Latinos is a collaborative study supported by contracts from the National Heart, Lung, and Blood Institute (NHLBI) to the University of North Carolina (HHSN268201300001I / N01-HC-65233), University of Miami (HHSN268201300004I / N01-HC-65234), Albert Einstein College of Medicine (HHSN268201300002I / N01-HC-65235), University of Illinois at Chicago (HHSN268201300003I / N01-HC-65236 Northwestern Univ), and San Diego State University (HHSN268201300005I / N01-HC-65237). The following Institutes/Centers/Offices have contributed to the HCHS/SOL through a transfer of funds to the NHLBI: National Institute on Minority Health and Health Disparities, National Institute on Deafness and Other Communication Disorders, National Institute of Dental and Craniofacial Research, National Institute of Diabetes and Digestive and Kidney Diseases, National Institute of Neurological Disorders and Stroke, NIH Institution-Office of Dietary Supplements.

Additional funding for the “Gut Origins of Latino Diabetes” (GOLD) ancillary study to HCHS/SOL was provided by 1R01MD011389-01 from the National Institute on Minority Health and Health Disparities.

The Genetic Analysis Center at the University of Washington was supported by NHLBI and NIDCR contracts (HHSN268201300005C AM03 and MOD03).

## Author information

### Author contributions

M.C.W. and R.C.K. oversaw the study. The methodology for the covariate-adjusted KRV was developed by M.C.W. and H.L., with contributions from W.L., X.H., X.Z. and N.Z. A.M.P. and H.L. implemented the covariate-adjusted KRV method into the R-based software. H.L. conducted simulations and analyzed the HCHS/SOL GOLD data using the covariate-adjusted KRV. R.C.K., R.K. and R.D.B. conceived of the HCHS/SOL GOLD study. R.C.K., Q.Q., R.K. and R.D.B. obtained funding for the HCHS/SOL GOLD study. R.C.K., Q.Q. and R.D.B. collected the data and specimens from the HCHS/SOL participants. R.D.B. performed the processing of the HCHS/SOL fecal samples. R.K. performed the gut microbial sequencing analysis for the HCHS/SOL GOLD study. X.H., J-Y.M. and J.S.W-N. performed pre-processing of the HCHS/SOL GOLD data. H.L. and M.C.W. drafted the manuscript, with contributions from W.L., X.H., J-Y.M., J.S.W-N., X.Z., A.M.P., N.Z., A.Z., R.K., Q.Q., R.D.B. and R.C.K.

### Correspondence

Correspondence and requests for materials should be addressed to M.C.W.

