## Supplementary Material for "Kernel-based genetic association analysis for microbiome phenotypes identifies host genetic drivers of beta-diversity"

### S.1 Supplementary tables and figures

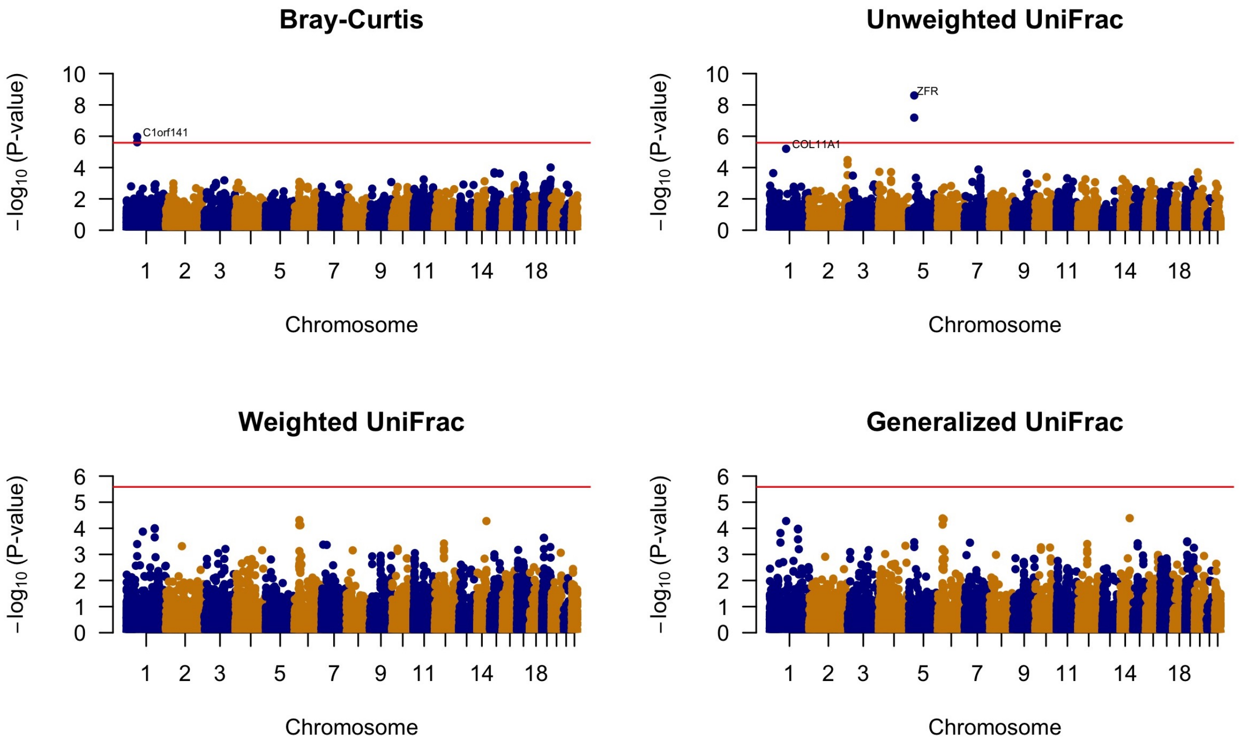


Figure S1: Manhattan plots from the first-stage (gene-level) analysis of the HCHS/SOL data, using the PC-adjusted KRV. Each plot corresponds to a distinct microbiome kernel. The top 5 PCs of genome-wide genetic variability were adjusted. The red lines represent the genome-wide significance threshold ($\alpha = 2.6 \times 10^{-6}$).

Table S1: P-values for the significant genes from Table 1 when additional covariates were adjusted in the KRV analysis.

| Microbiome kernel | Genes | Number of common variants | P-value |
|---|---|---|---|
| Bray-Curtis | *C1orf141* | 484 | $2.3 \times 10^{-5}$ |
| | *IL23R* | 284 | $3.2 \times 10^{-5}$ |
| Unweighted UniFrac | *MTMR12* | 174 | $2.1 \times 10^{-7}$ |
| | *ZFR* | 288 | $2.5 \times 10^{-8}$ |

Adjusted covariates include the top 5 PCs of genome-wide genetic variability, age, gender and study sites. The analysis was performed on 1096 unrelated individuals where all relevant data were available.

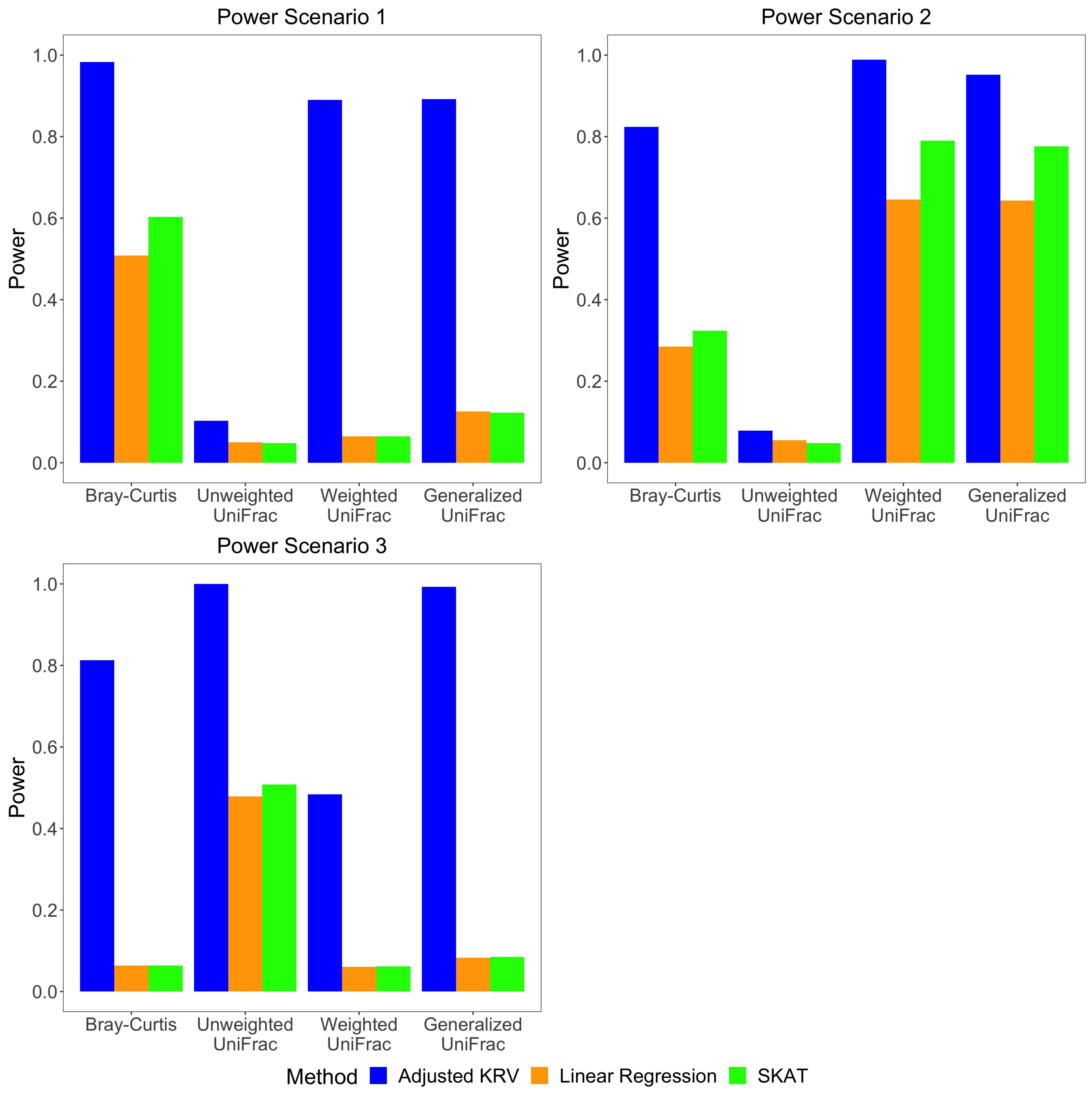


Figure S2: Empirical power of covariate-adjusted KRV and competing methods at nominal level $\alpha = 0.05$ for different microbiome kernels under large effect sizes. Linear kernel was used for genetic data.
